# Repression of CENP-A assembly in metaphase requires HJURP phosphorylation and inhibition by M18BP1

**DOI:** 10.1101/2021.10.28.466278

**Authors:** Julio C. Flores Servin, Aaron F. Straight

## Abstract

Centromeres are the foundation for mitotic kinetochore assembly and thus are essential for chromosome segregation. Centromeres are epigenetically defined by nucleosomes containing the histone H3 variant CENP-A. CENP-A nucleosome assembly is uncoupled from replication and occurs in G1 but how cells control this timing is incompletely understood. The formation of CENP-A nucleosomes in vertebrates requires CENP-C and the Mis18 complex which recruit the CENP-A chaperone HJURP to centromeres. Using a cell-free system for centromere assembly in *X. laevis* egg extracts, we discover two activities that inhibit CENP-A assembly in metaphase. HJURP phosphorylation prevents the interaction between HJURP and CENP-C in metaphase, blocking the delivery of soluble CENP-A to centromeres. Non-phosphorylatable mutants of HJURP constitutively bind CENP-C in metaphase but are not sufficient for new CENP-A assembly. We find that the M18BP1.S subunit of the Mis18 complex also binds to CENP-C to competitively inhibit HJURP’s access to centromeres. Removal of these two inhibitory activities causes CENP-A assembly in metaphase.

**SUMMARY:** Vertebrate CENP-A assembly is normally restricted to G1 phase. Two inhibitory activities, phosphorylation of HJURP and competitive binding of M18BP1.S to CENP-C, block HJURP’s access to the metaphase centromere. Removal of these inhibitory activities causes CENP-A assembly in metaphase.

## INTRODUCTION

Faithful chromosome segregation during mitosis requires that each pair of sister chromatids attach to microtubules from opposite poles of the mitotic spindle, via a chromosomal structure termed the kinetochore. During mitosis, the kinetochore assembles on a specialized chromatin region called the centromere. In many eukaryotes, functional centromeres are epigenetically defined by the presence of the histone H3 variant, centromere protein A (CENP-A) (Warburton et al., 1997; Palmer et al., 1987; Sullivan et al., 1994; Earnshaw and Rothfield, 1985; Palmer et al., 1991; Meluh et al., 1998; Takahashi et al., 2000; Henikoff et al., 2000). Perturbation or loss of CENP-A results in the loss of centromeres and kinetochores, chromosome missegregation, aneuploidy, and cell death (Stoler et al., 1995; Blower and Karpen, 2001; Howman et al., 2000; Régnier et al., 2005; Goshima et al., 2003). Furthermore, ectopic incorporation of CENP-A at non-centromeric regions can lead to neocentromere formation, chromosome instability, and breakage events (Heun et al., 2006; Barnhart et al., 2011; Hori et al., 2013). Therefore, accurate chromosome segregation requires proper CENP-A nucleosome formation for centromere maintenance.

During DNA replication nucleosomes are distributed between newly replicated chromatids and new histone H3 nucleosome assembly is coupled to passage of the replication fork to regenerate chromatin (Groth et al., 2007; Ramachandran and Henikoff, 2015). Unlike histone H3 nucleosomes, the timing of new CENP-A assembly is uncoupled from DNA replication in many organisms. In *Arabidopsis thaliana* and *Schizosaccharomyces pombe* new CENP-A assembly occurs in G2 (Lermontova et al., 2006; Lando et al., 2012; Takayama et al., 2008) and in *Drosophila melanogaster* during M-phase in somatic cells (Mellone et al., 2011) and anaphase in early embryos (Schuh et al., 2007). In vertebrates and *Caenorhabditis elegans*, CENP-A nucleosomes are assembled after mitotic exit in G1 (Jansen et al., 2007; Maddox et al., 2007; Moree et al., 2011; Bernad et al., 2011; Silva et al., 2012). However, we do not fully understand the importance of uncoupling CENP-A assembly from DNA replication or how cells control the timing of CENP-A nucleosome formation.

Three factors play essential roles in the assembly of new CENP-A in vertebrates. The CENP-A specific histone chaperone Holliday junction recognition protein (HJURP) forms a complex with a heterodimer of CENP-A and histone H4 to deliver new CENP-A to the centromere (Dunleavy et al., 2009; Foltz et al., 2009). The hetero-octameric Mis18 complex of three proteins (Mis18α(2):Mis18β(4):M18BP1/KNL2(2)) is required for localizing the HJURP chaperone complex to centromeres in G1 (Fujita et al., 2007; Maddox et al., 2007; Moree et al., 2011; Pan et al., 2019). The targeting of HJURP to centromeres through the Mis18 complex involves two conserved repeat domains in HJURP that directly bind to the Mis18 complex through interactions with Mis18αβ (Nardi et al., 2016; Pan et al., 2017). Finally, centromere protein C (CENP-C) binds directly to CENP-A nucleosomes (Carroll et al., 2010) and also binds HJURP and the Mis18 complex. HJURP (Tachiwana et al., 2015; French et al., 2017), the M18BP1 protein (Moree et al., 2011; Dambacher et al., 2012), and the Mis18β subunit (Stellfox et al., 2016) of the Mis18 complex all interact with the C-terminal domain of CENP-C. Thus, the coordinated interactions between HJURP, the Mis18 complex, and CENP-C govern the timing and localization of CENP-A assembly at centromeres.

Vertebrate CENP-C binds to centromeres throughout the cell cycle thus changes in its localization are not thought to regulate the timing of CENP-A assembly. In humans, the Mis18 complex and HJURP associate with centromeres exclusively in G1 thus it has been proposed that regulation of their localization controls the G1 assembly of CENP-A (Fujita et al., 2007; Maddox et al., 2007; Dunleavy et al., 2009; Foltz et al., 2009). However, in chicken DT-40 cells and *Xenopus laevis* egg extracts, M18BP1 constitutively binds to centromeres (Moree et al., 2011; Hori et al., 2017; French et al., 2017). One common feature between these systems is that the G1 timing of new CENP-A assembly is controlled by mitotic kinases that disrupt the interactions between the Mis18 complex, HJURP and CENP-C (Silva et al., 2012; Wang et al., 2014; Müller et al., 2014). Cyclin dependent kinase 1 (CDK1) activity negatively regulates the centromere localization of M18BP1 in humans until cyclin degradation and exit from mitosis (Silva et al., 2012; Jansen et al., 2007; McKinley and Cheeseman, 2014; Stankovic et al., 2017). CDK1 phosphorylation of M18BP1 drives the disassembly of the Mis18 complex by disrupting the interaction between M18BP1 and Mis18α/β (McKinley and Cheeseman, 2014; Pan et al., 2017; Spiller et al., 2017; French and Straight, 2019). In addition, CDK1 phosphorylation of HJURP disrupts HJURP localization to centromeres (Müller et al., 2014; Stankovic et al., 2017) by disrupting both HJURP/Mis18 complex interaction (Wang et al., 2014; Stellfox et al., 2016; French et al., 2017; Pan et al., 2019) and HJURP/CENP-C interaction (French et al., 2017) but the mechanisms for how CDK1 activity controls these interactions is not fully understood.

The constitutive localization of *Xenopus laevis* M18BP1 to centromeres is explained in part by a conserved domain present in the frog, chicken, and plant M18BP1 proteins, but absent in mammals, that directly binds to CENP-A nucleosomes using the same mechanism as CENP-C (Hori et al., 2017; French et al., 2017; Sandmann et al., 2017; Kral, 2015). *Xenopus laevis* is allotetraploid, containing two related subgenomes. As a result, *X. laevis* has two homeologs of M18BP1 termed M18BP1.S and M18BP1.L (previously termed M18BP1-1 and M18BP1-2 respectively (Moree et al., 2011)). Both the S and L isoforms bind to centromeres throughout interphase using direct binding to CENP-A nucleosomes. However in metaphase, Cdk phosphorylation causes a switch in the mechanism of M18BP1 targeting, inhibiting interaction with the CENP-A nucleosomes and promoting the interaction of M18BP1.S with CENP-C (Moree et al., 2011; French et al., 2017; French and Straight, 2019).

In this work we take advantage of the *Xenopus laevis* system to uncover new mechanisms controlling HJURP centromere targeting in vertebrates. We find that mitotic phosphorylation of HJURP on serine 220 prevents its interaction with the cupin dimerization domain of CENP-C in metaphase. Phosphomimetic mutants of HJURP S220 or CENP-C mutants that disrupt CENP-C dimerization prevent HJURP from localizing to centromeres in interphase and inhibit new CENP-A assembly. Non-phosphorylatable mutants of HJURP result in constitutive interaction between HJURP and CENP-C in metaphase, yet do not localize to centromeres or cause premature CENP-A assembly. We find that M18BP1.S provides a second inhibitory mechanism preventing premature CENP-A assembly in metaphase by binding to the CENP-C cupin domain and preventing HJURP from localizing to centromeres. When we remove these two negative regulators, mitotic phosphorylation of HJURP and competitive inhibition by M18BP1.S, HJURP localizes to centromeres and drives new CENP-A assembly in metaphase. Our studies identify dual regulatory mechanisms controlling the cell cycle specific assembly of CENP-A.

## RESULTS

### CENP-A assembly requires HJURP binding to the CENP-C cupin domain

HJURP binds directly to CENP-C in interphase and this interaction is required for new CENP-A assembly (Tachiwana et al., 2015; French et al., 2017). To better understand the determinants of this interaction, we measured the binding of HJURP to a series of non-overlapping CENP-C truncations (Fig. 1a). We translated full-length or truncated Myc-tagged CENP-C in rabbit reticulocyte lysates and added the translated proteins to *X. laevis* egg extracts that had been depleted of the endogenous CENP-C. After immunoprecipitation of Myc-CENP-C and western blotting for HJURP we observed interphase-specific association of HJURP with full-length CENP-C and a C-terminal truncation of CENP-C containing the cupin domain (Fig. 1b). To test if CENP-C dimerization is required for interaction with HJURP we created CENP-C mutants that should fail to dimerize (F1332A and F1393A) based on previously reported structural and homology data (Cohen et al., 2008; Carroll et al., 2010; Medina-Pritchard et al., 2020) and confirmed that these mutants disrupted CENP-C dimerization in egg extracts (Extended Data Fig. 1a). We tested if CENP-C dimerization mutants interacted with HJURP in interphase extracts and found that the CENP-C dimerization mutants, but not mutants in the CENP-A nucleosome binding domains, inhibited HJURP/CENP-C interaction (Fig. 1c). Together, these results indicate that the CENP-C C-terminus and CENP-C dimerization are required for proper HJURP association.

**Figure 1.**
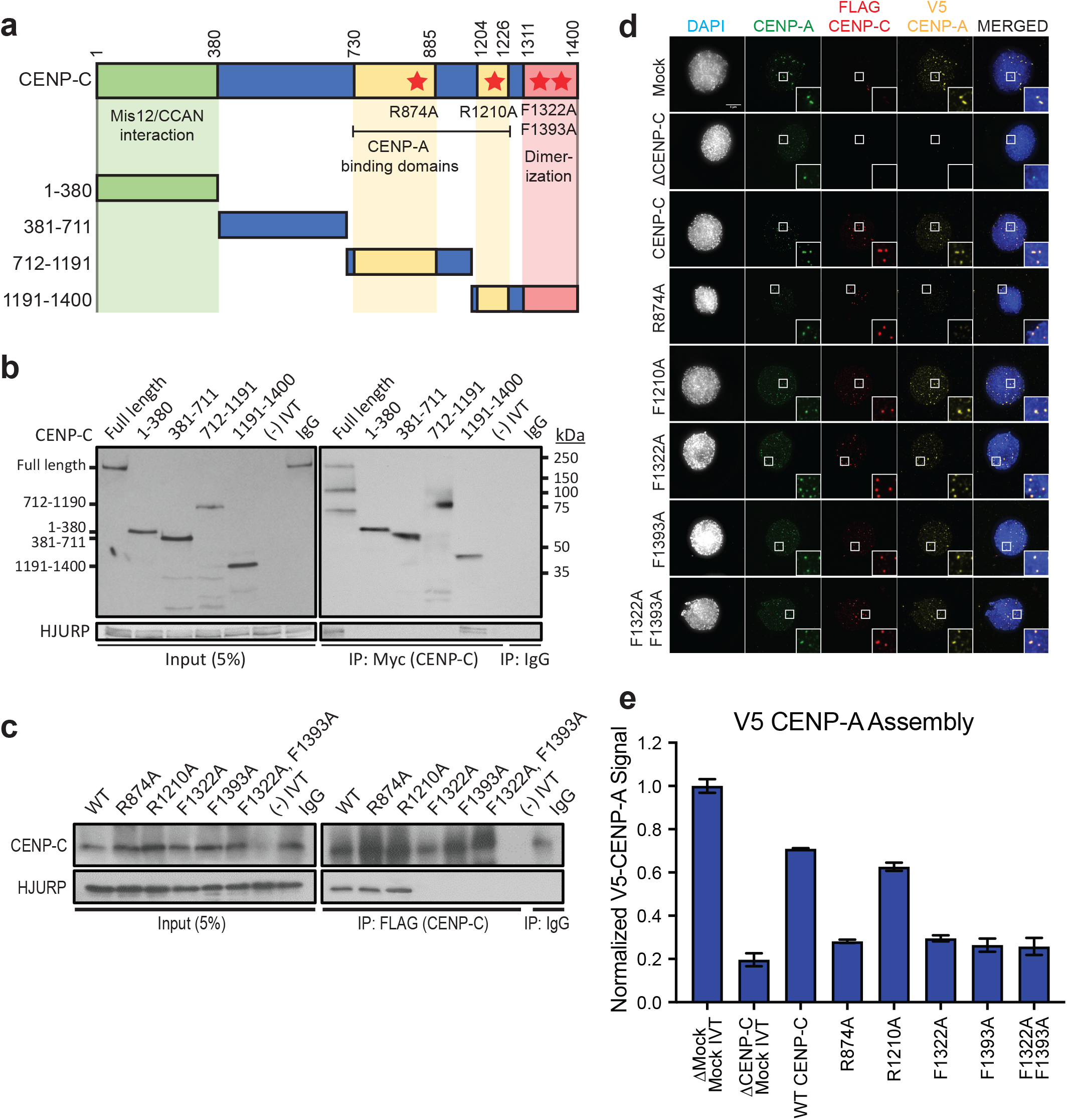
HJURP interaction with the CENP-C cupin domain is required for CENP-A assembly. a) Schematic of CENP-C truncations and mutations. The CENP-C amino acid residue numbers for each truncation are listed to the left. The domains of CENP-C that interact with Mis12/CCAN, interact with CENP-A nucleosomes, and the cupin dimerization domain are highlighted in green, yellow and red respectively. The residue numbers of the boundaries of each domain are listed on top. Mutations that inhibit CENP-A nucleosome binding and dimerization are highlighted with red stars. b) The C-terminus of CENP-C binds to HJURP. Interphase extract depleted of endogenous CENP-C was supplemented with Myc-CENP-C truncations. Co-immunoprecipitation of HJURP was assayed by anti-HJURP immunoblotting after immunoprecipitation of CENP-C truncations from interphase *X. laevis* egg extracts. The left panels show 5% of the input used in the immunoprecipitation and the right panels show the CENP-C (top panels) and HJURP (bottom panels) in the immunoprecipitates. Each fragment or control (-IVT: scrambled DNA translated *in-vitro*, IgG: nonspecific mouse IgG antibody) is listed above the panels, the size of each fragment is shown to the left and the molecular weight to the right. c) CENP-C cupin domain mutations affect HJURP interaction. Interphase extract depleted of endogenous CENP-C was supplemented with the specified FLAG-CENP-C mutants. Co-immunoprecipitation of endogenous HJURP with each FLAG-CENP-C protein was assayed by anti-HJURP immunoblotting (bottom panels). The levels of CENP-C in the extract and immunoprecipitation are shown in the top panels. Mock precipitations using scrambled DNA translated *in-vitro* (-IVT) or whole mouse IgG served are indicated. The left panels contain 5% of the input material and the right panels contain the immunoprecipitations. d) CENP-C cupin domain mutations disrupt CENP-A assembly. Representative images of sperm nuclei incubated in CENP-C depleted interphase *Xenopus* egg extracts complemented with the CENP-C mutant indicated. Extracts were supplemented with RNA encoding V5-CENP-A and *in vitro* translated HJURP protein to assay CENP-A assembly. Immuno-localized protein is specified above. Scale bar, 5 μm. Insets are magnified 300%. e) Quantification of V5-CENP-A assembly shown in **(d)**. Values are normalized to unsupplemented, mock-depleted extract. Plot shows mean V5-CENP-A signal on sperm chromatin ± SEM of at least three experiments.

CENP-C binding to CENP-A nucleosomes and CENP-C dimerization are required for proper localization of CENP-C to centromeres (Carroll et al., 2010). We tested whether CENP-C binding to CENP-A nucleosomes or dimerization were required for centromeric HJURP localization and new CENP-A assembly. We complemented CENP-C depleted interphase extracts with CENP-C dimerization and CENP-A nucleosome binding mutants and measured the centromeric levels of HJURP and the assembly of new CENP-A. Extracts rescued with the CENP-A binding deficient mutant CENP-C^R874A^ had a 72% ± 3% reduction in CENP-A assembly, compared to wild-type. Extracts complemented with CENP-C dimerization mutants CENP-C^F1332A^, CENP-C^F1393A^, and CENP-C^F1332A/F1393A^, showed a 76% ± 9%, 74% ± 6%, and 70% ± 4% reduction (respectively) in new CENP-A assembly compared to wild type (Fig. 1d,e, and Extended Data Fig. 1b). Both HJURP and CENP-C localization to centromeres were reduced in extracts complemented with CENP-C dimerization mutants (Extended Data Fig. 1c,d). These data indicate that *X. laevis* CENP-C dimerization is necessary for HJURP binding to CENP-C, HJURP centromere localization, and new CENP-A assembly.

### HJURP interaction with CENP-C in interphase is controlled by a conserved mitotic phosphorylation site

To map the binding domain in HJURP that interacts with CENP-C we measured the binding between a series of FLAG-HJURP truncations and CENP-C in interphase *Xenopus laevis* egg extracts (Fig. 2a). We translated full-length or truncated FLAG-tagged HJURP in rabbit reticulocyte extracts, added the translated protein to interphase extracts, and assessed binding to CENP-C by immunoprecipitation. We found that the N-terminus of HJURP from amino acids 1–225 was sufficient for CENP-C interaction but that the interaction was lost when HJURP was truncated to amino acids 1–205 (Fig. 2b,c). HJURP truncations encompassing amino acids 201 to the C-terminus were unable to bind CENP-C (Fig. 2b and Extended Data Fig. 2a,b). These data indicate that the N-terminal 225 amino acids of HJURP is sufficient for its interaction with CENP-C and residues 205-225 are essential for this interaction.

**Figure 2.**
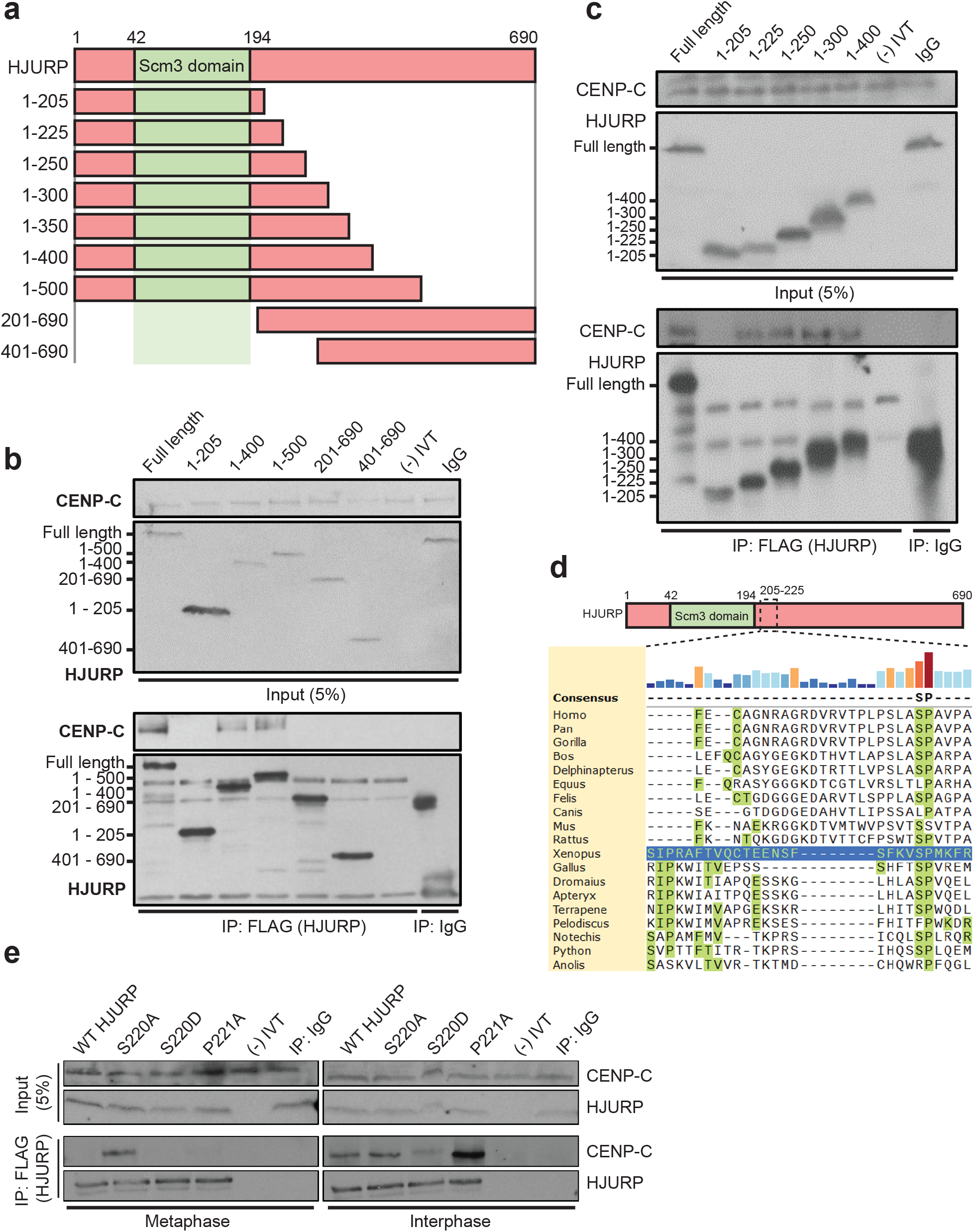
A conserved SP site on the HJURP N-terminus regulates its interaction with CENP-C. a) Schematic representation of HJURP truncations used to identify the CENP-C interacting region in **(b)** and **(c)**. b) A fragment of HJURP spanning amino acids 1–400 binds to CENP-C. Interphase extract was supplemented with the FLAG-HJURP truncations specified at the top of the panel. Following FLAG-HJURP immunoprecipitation, co-precipitation of CENP-C was assayed by anti-CENP-C immunoblotting. For (b,c, and e) mock precipitations using scrambled DNA translated *in-vitro* (-IVT) or whole mouse IgG serve as negative controls. For **(b)** and **(c)** the top panel shows 5% of the input material, the bottom panel shows the immunoprecipitates with the sizes of HJURP fragments indicated on the left. c) HJURP amino acid residues 205–225 are required for its interaction with CENP-C. Interphase extract was supplemented with the specified FLAG-HJURP truncations. Co-immunoprecipitation of endogenous CENP-C was assayed by anti-CENP-C immunoblotting following FLAG-HJURP immunoprecipitation. d) HJURP contains a conserved N-terminal serine/proline site across most vertebrates. HJURP sequences are aligned using *X. laevis* HJURP^205-225^ as a reference (blue row). e) HJURP serine 220 (HJURP^S220^) regulates association with CENP-C. Interphase and metaphase extracts were supplemented with the indicated FLAG-HJURP mutants. Co-immunoprecipitation of endogenous CENP-C was assayed by anti-CENP-C immunoblotting following FLAG-HJURP precipitation. The top panel shows 5% of the input material, the bottom panel shows the immunoprecipitates.

We aligned *Xenopus* HJURP 205-225 with diverse HJURP orthologs which revealed a single highly conserved serine (S220) and proline (P221) motif within this CENP-C recognition region (Fig. 2d). The core consensus site for mitotic phosphorylation by cyclin dependent kinases on serine or threonine is S/T-P-X-K/R (Holt et al., 2009) and the conserved region we find in the *X. laevis* HJURP is S-P-M-K, suggesting that this SP site may be the target of cyclin dependent mitotic kinase activity. In humans, the homologous site has previously been identified as a mitotically phosphorylated residue in phosphoproteomic studies of HJURP (Stankovic et al., 2017) thus we hypothesized that cell cycle-dependent phosphorylation of the conserved serine 220 might regulate the interaction of *Xenopus* HJURP with CENP-C.

### HJURP S220 and P221 regulate HJURP’s interaction with CENP-C

To test whether the conserved SP220/221 site on HJURP regulates its interaction with CENP-C, we generated mutations in *Xenopus* FLAG-HJURP in which either the S220 or P221 residues were mutated to alanine to prevent phosphorylation or the S220 residue was mutated to aspartic acid to mimic phosphorylation. We expressed these mutants in metaphase and interphase *Xenopus laevis* egg extracts followed by immunoprecipitation of the HJURP mutants and western blotting for CENP-C. Surprisingly, we found that in metaphase extracts the nonphosphorylatable HJURP^S220A^ mutant interacted with CENP-C at levels similar to that of wild type HJURP in interphase (Fig. 2e). In interphase, the HJURP^S220A^ mutant exhibited no difference in its interaction with CENP-C compared to wild-type HJURP. Remarkably, in interphase extracts, HJURP^P221A^ bound significantly more CENP-C, suggesting proline 221 plays an important role in CENP-C interaction (Fig. 2e). In contrast, the phosphomimetic HJURP^S220D^ mutant showed significant reduction in CENP-C interaction during interphase (Fig. 22). When we expressed the phosphomimetic mutant of HJURP and CENP-C in reticulocyte extracts without incubation in *X. laevis* egg extract we did not observe inhibition of binding between the two proteins. This suggests that *Xenopus* egg extracts contain an activity that is required for the inhibition of interaction between HJURP and CENP-C (Extended Data Fig. 2c). Mutation of S220 and P221 had no effect on HJURP interaction with CENP-A/H4 dimer (Extended Data Fig. 2d,e). Together these data suggest that metaphase phosphorylation of HJURP on S220 prevents its interaction with CENP-C until CDK activity declines in interphase, thereby restricting the timing of new CENP-A assembly to interphase.

### Mutations in HJURP^S220/P221^ disrupt HJURP centromere localization and new CENP-A assembly

We tested whether HJURP S220 phosphorylation controls HJURP localization to sperm chromatin centromeres and new CENP-A assembly by measuring the levels of wild type and S220/P221 mutant HJURP at centromeres and the levels of new CENP-A assembly in metaphase and interphase extracts. In metaphase, we observed no localization of wild type or mutant HJURP to sperm chromatin centromeres even though the S220A and P221A mutations bind CENP-C in metaphase (Fig. 3a,b). Consistent with the lack of localization of HJURP in metaphase we observed no new CENP-A assembly in wild type or HJURP mutants (Fig. 3c). In interphase extracts, HJURP^S220D^ showed a 70% ± 4% reduction of centromere localization (Fig. 3a,b) and a 75% ± 3% reduction in new CENP-A assembly (Fig. 3c), consistent with an inhibitory role for S220 phosphorylation on CENP-C interaction. HJURP^S220A^ demonstrated a slight increase in localization to centromeres while HJURP^P221A^ showed an increase of 180% ± 40% in centromere localization (Fig. 3a,b). HJURP^S220A^ and HJURP^P221A^ exhibited an increase of 37% ± 6% and 117% ± 12% in new CENP-A assembly respectively (Fig. 3c), compared to the wild type control, consistent with the increased localization and binding of the HJURP^S220A^ and HJURP^P221A^ mutants to CENP-C (Fig. 2e).

**Figure 3.**
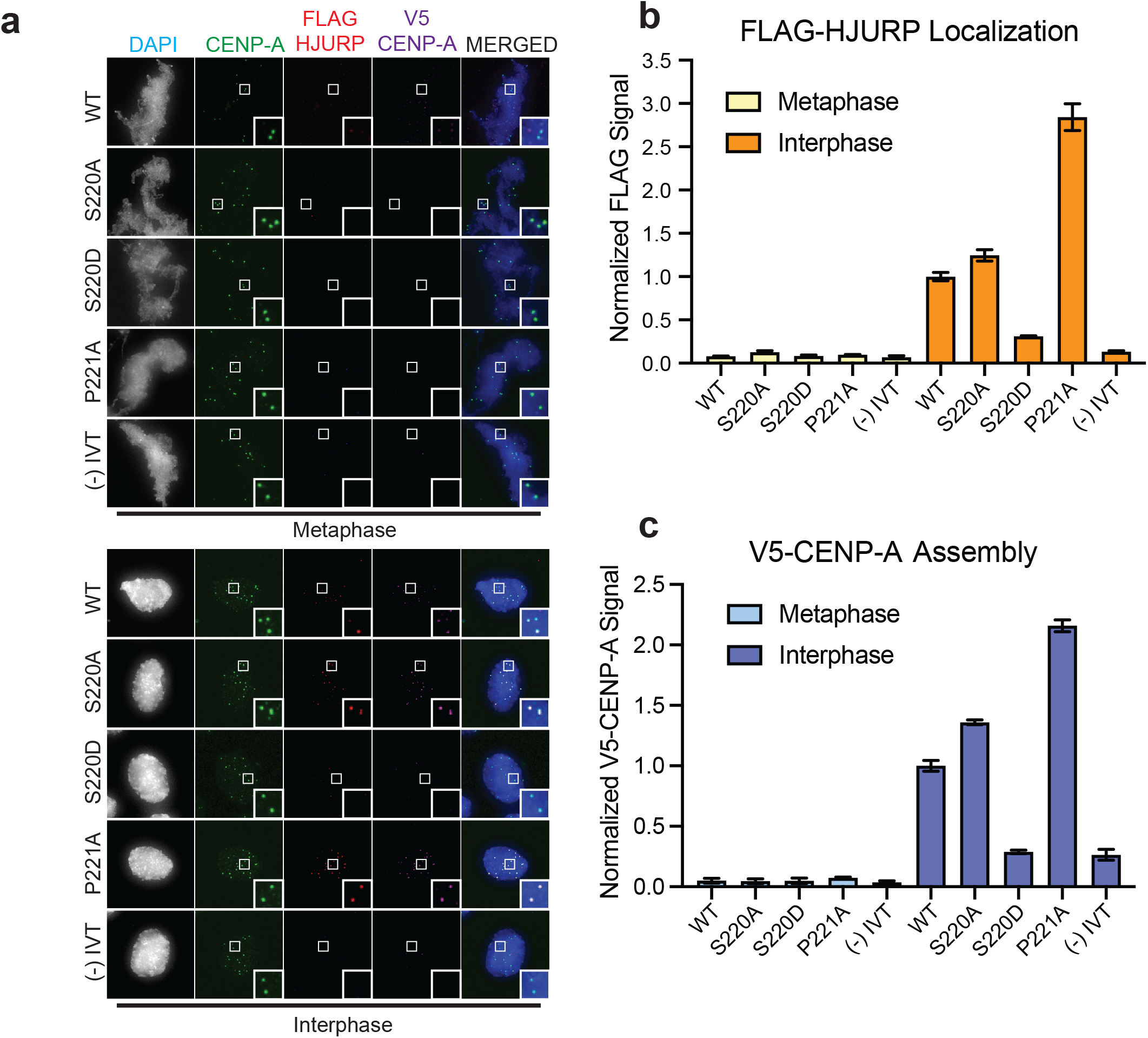
HJURP^S220^ regulates CENP-A assembly. a) HJURP containing the phosphomimetic S220D mutation inhibits new CENP-A assembly on sperm chromatin. Representative images of sperm nuclei incubated with metaphase or interphase *Xenopus* egg extracts were complemented with the indicated HJURP mutant. Metaphase and interphase extracts were complemented with RNA encoding V5-CENP-A to assay for new CENP-A assembly. Immuno-localized protein is specified above. Insets are magnified 300%. b) Quantification of the levels of FLAG-HJURP at centromeres from **(a)**. Values are normalized to wild type FLAG-HJURP signal in interphase extract. The plot shows the mean FLAG-HJURP signal on sperm chromatin ± SEM of at least three experiments. c) Quantification of V5-CENP-A assembly from **(a)**. Values are normalized to V5-CENP-A assembly in wild type HJURP condition in interphase extracts. The plot shows the mean V5-CENP-A assembly signal on sperm chromatin ± SEM of at least three experiments.

Together, these results indicate that HJURP^S220^ phosphorylation prevents interaction with CENP-C in metaphase and thus inhibits HJURP localization and new CENP-A assembly. However, HJURP^S220A^, despite being able to interact with CENP-C during metaphase, is unable to promote HJURP localization or new CENP-A assembly. This suggests that an additional inhibitory mechanism prevents premature HJURP localization and CENP-A assembly in metaphase.

### M18BP1.S prevents HJURP localization and CENP-A assembly during metaphase

During metaphase, M18BP1.S interacts with the C-terminus of CENP-C in the same region where HJURP binds to CENP-C (French and Straight, 2019)). To test whether M18BP1.S competitively inhibits HJURP binding to CENP-C, we depleted M18BP1.S and M18BP1.L from *X. laevis* egg extracts (Extended Data Fig. 3a) and measured the levels of wild type and S220/P221 mutant FLAG-HJURP localization to sperm centromeres. In mock depleted metaphase extracts, where M18BP1 was still present, we observed no change in centromeric HJURP localization (Fig. 4b,d). However, when M18BP1.S and M18BP1.L were depleted, the levels of HJURP^S220A^ and HJURP^P221A^ at metaphase centromeres rose to 23% ± 4% and 50% ± 7% compared to mock depleted interphase extracts (Fig. 4a,b). Complementation of the M18BP1.S/M18BP1.L depleted extracts with M18BP1.S suppressed the localization of HJURP^S220A^ and HJURP^P221A^ to 11% ± 4% and 14% ± 8%, compared to wild type HJURP levels in mock depleted interphase extracts (Fig. 4a,b). These observations suggest that the presence of M18BP1.S in metaphase extracts inhibits the localization of HJURP to centromeres.

**Figure 4.**
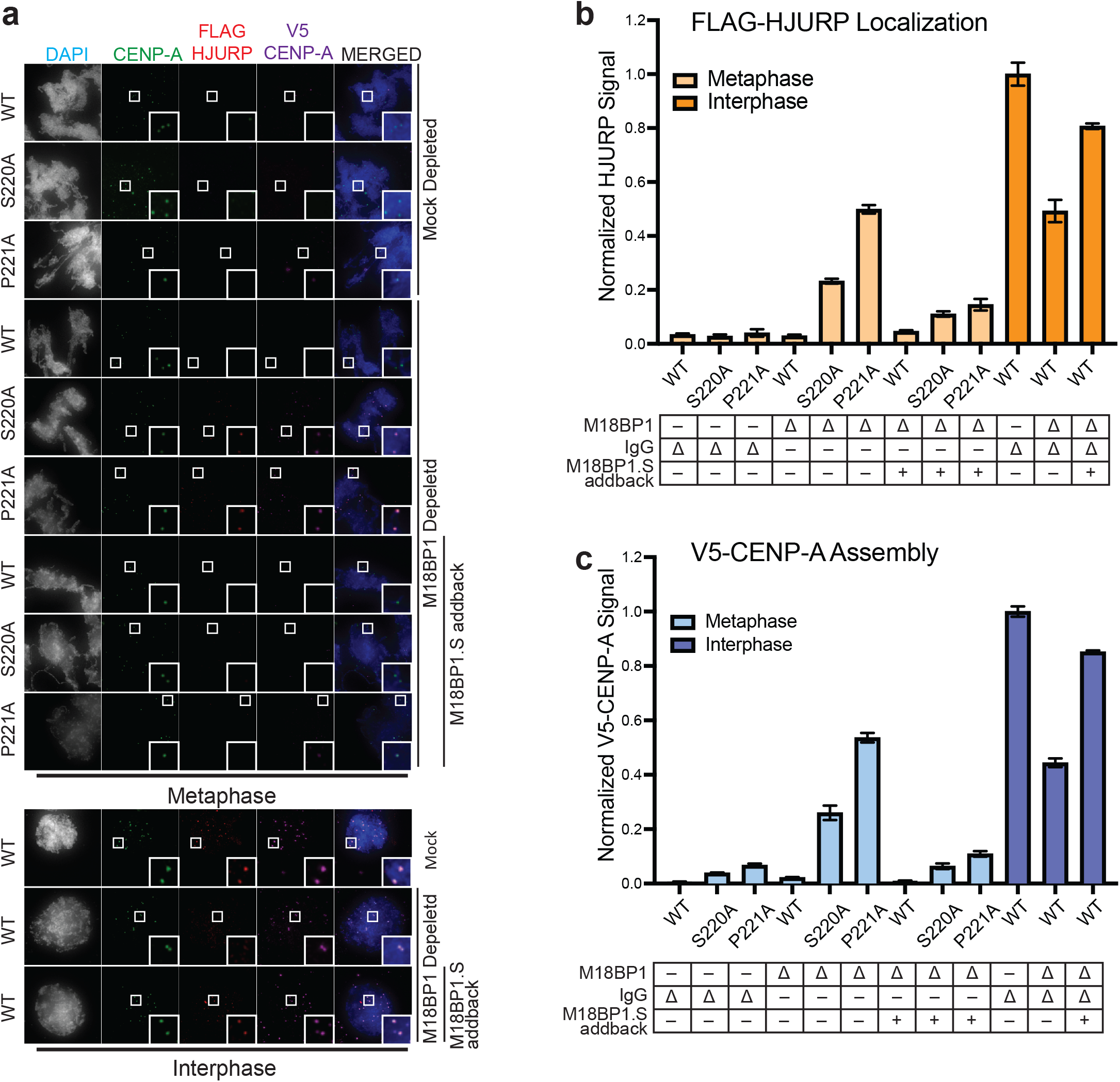
M18BP1.S prevents HJURP association with CENP-C and new CENP-A assembly during metaphase. a) M18BP1 depletion causes premature HJURP^S220A^ centromere localization in metaphase and new CENP-A assembly. Representative images of sperm nuclei incubated with M18BP1-depleted metaphase or interphase *Xenopus* egg extracts complemented with the HJURP mutants indicated to the left of the images. Right hand labels indicate mock depleted or M18BP1 depleted and M18BP1.S add-back for each condition. Metaphase and interphase extracts were supplemented with RNA encoding V5-CENP-A to assay for new CENP-A assembly. Immuno-localized protein is specified above. Insets are magnified 300%. b) Quantification of FLAG-HJURP centromere localization in **(a)**. Values are normalized to the wild type FLAG-HJURP centromere signal in mock depleted interphase extract. Bottom rows indicate antibody depletion (Δ) (M18BP1 or IgG antibody) and M18BP1.S add-back (+) for each condition. Plot shows mean FLAG-HJURP signal on sperm chromatin ± SEM of at least three experiments. c) Quantification of V5-CENP-A assembly in **(a)**. Values are normalized to the V5-CENP-A assembly signal in wild type HJURP condition in mock depleted interphase extracts. Bottom rows indicate depletion (Δ) status (M18BP1 or IgG antibody) and M18BP1.S add-back (+) for each condition. Plot shows mean V5-CENP-A assembly signal on sperm chromatin ± SEM of at least three experiments.

We tested whether premature localization of mutant HJURP in metaphase was sufficient to promote new CENP-A assembly on sperm centromeres. In mock depleted extracts neither wild type nor mutant HJURP promoted new CENP-A assembly (Fig. 4a,c). However, addition of HJURP^S220A^ and HJURP^P221A^ to M18BP1.S and M18BP1.L depleted metaphase extracts drove new CENP-A assembly to 26% ± 12% and 53% ± 9% higher than extracts complemented with wild type HJURP (Fig. 4a,c). Complementing these extracts with M18BP1.S inhibited new CENP-A assembly to levels similar to mock depleted conditions (Fig. 4c). This inhibition was specific to M18BP1.S as only M18BP1.S and not M18BP1.L or a mutant in M18BP1.S that cannot bind CENP-C or localize to centromeres, was able to prevent premature HJURP localization and new CENP-A assembly (Extended Data Fig. 3b,c,d,e). Depletion of M18BP1 in interphase reduced CENP-A assembly to 44% ± 2% of the level of mock depleted interphase extracts, demonstrating that although M18BP1.S acts as an inhibitor of CENP-A assembly in metaphase, full CENP-A assembly requires the presence of M18BP1 in interphase (Fig. 4a,c). Co-depletion of CENP-C in these experiments prevented metaphase wild type and mutant HJURP localization and new CENP-A assembly, demonstrating the dependence of CENP-A assembly on CENP-C (Extended Data Fig. 4). Consistent with the role of CENP-C in mediating the effect of HJURP mutants on new CENP-A assembly, depletion of CENP-C from interphase extracts reduced the effects of the HJURP mutants to the level of the wild type protein (Extended Data Fig. 4). Thus, M18BP1.S acts to inhibit HJURP localization and new CENP-A assembly in metaphase by competing for the HJURP binding site on CENP-C.

We tested if addition of recombinant purified M18BP1.S^161–580^, the CENP-C interacting region on M18BP1.S (Fig. 5a), was sufficient to compete with HJURP for binding to CENP-C in metaphase egg extracts. Addition of MBP-6xHis tagged M18BP1.S^161–580^ (Fig. 5b) to mock and M18BP1 depleted metaphase arrested egg extracts reduced the interaction of HJURP^S220A^ and HJURP^P221A^ with CENP-C (Fig. 5c). Doubling the amount of M18BP1.S^161–580^ reduced the amount of CENP-C coprecipitated with HJURP to levels similar to those observed in mock depleted conditions (Fig. 5d). Mutations to the SANTA domain of M18BP1.S prevent its interaction with CENP-C in metaphase (French and Straight, 2019), thus we tested if addition of M18BP1.S^161–580^, containing previously defined SANTA mutations (French and Straight, 2019), would fail to compete with the HJURP and CENP-C interaction in M18BP1 depleted metaphase extracts. Addition of M18BP1.S^SANTA/161–580^ to M18BP1 depleted metaphase extracts partially reduced the ability of HJURP^S220A^ and HJURP^P221A^ to interact with CENP-C, suggesting that M18BP1.S competes for CENP-C binding to HJURP through its SANTA domain (Fig. 5d). Thus, M18BP1.S inhibits HJURP localization and new CENP-A assembly in metaphase by competing for the HJURP binding site on CENP-C (Fig. 6a).

**Figure 5.**
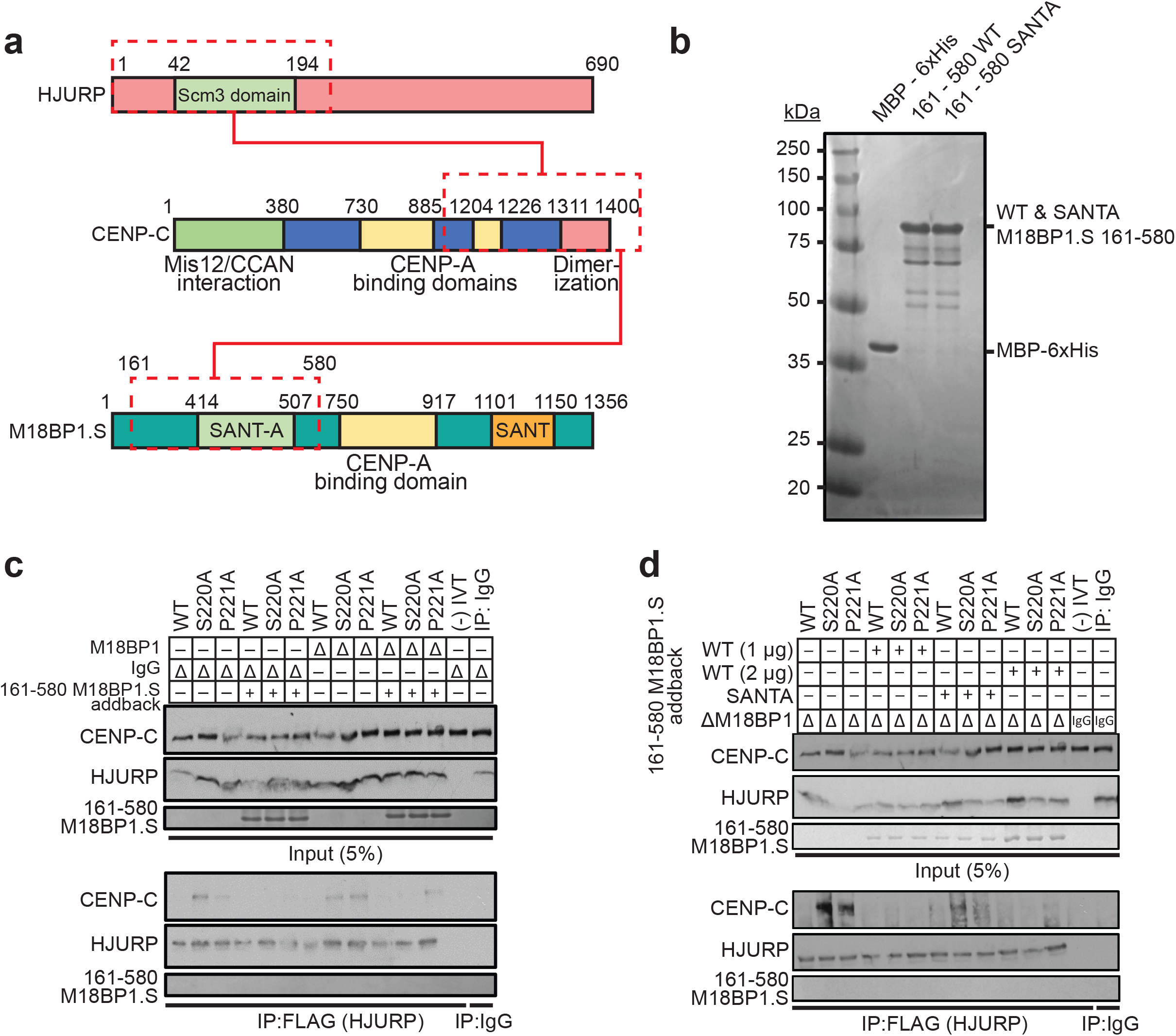
M18BP1.S competes for HJURP binding to CENP-C. a) Schematic of the domains of HJURP and M18BP1.S that compete for binding to the CENP-C C-terminus. b) Coomassie stained gel of purified proteins used in **(c)** and **(d)**. Proteins are indicated on top and their full length migration position to the right. The molecular weight standards in kDa are indicated to the left. c) Addition of M18BP1.S to undepleted or M18BP1 depleted metaphase extract competes for the interaction between HJURP^S220A^ or HJURP^P221A^ and CENP-C. HJURP mutants indicated across the top were added to metaphase *X. laevis* extract that had been depleted with M18BP1 antibody or mock depleted with IgG (Δ). Extract supplemented with the 151-580 fragment of M18BP1.S are indicated (+). The top panel contains 5% of the input material and the bottom panel contains the immunoprecipitates after FLAG-HJURP precipitation and western blotting for CENP-C, FLAG (HJURP) and M18BP1. d) The SANTA domain mutant of M18BP1.S that cannot bind CENP-C fails to compete for HJURP^S220A^ or HJURP^P221A^ binding. Extracts were manipulated as in **(c)** with the addition of two different concentrations of M18BP1.S^161-580^ or the addition of the M18BP1.S^SANTA^ mutant as indicated on the left. The top panel contains 5% of the input material and the bottom panel contains the immunoprecipitates after FLAG-HJURP precipitation and western blotting for CENP-C, FLAG (HJURP) and M18BP1. Mock precipitations using scrambled DNA translated *in-vitro* (-IVT) or whole mouse IgG serve as negative controls.

**Figure 6.**
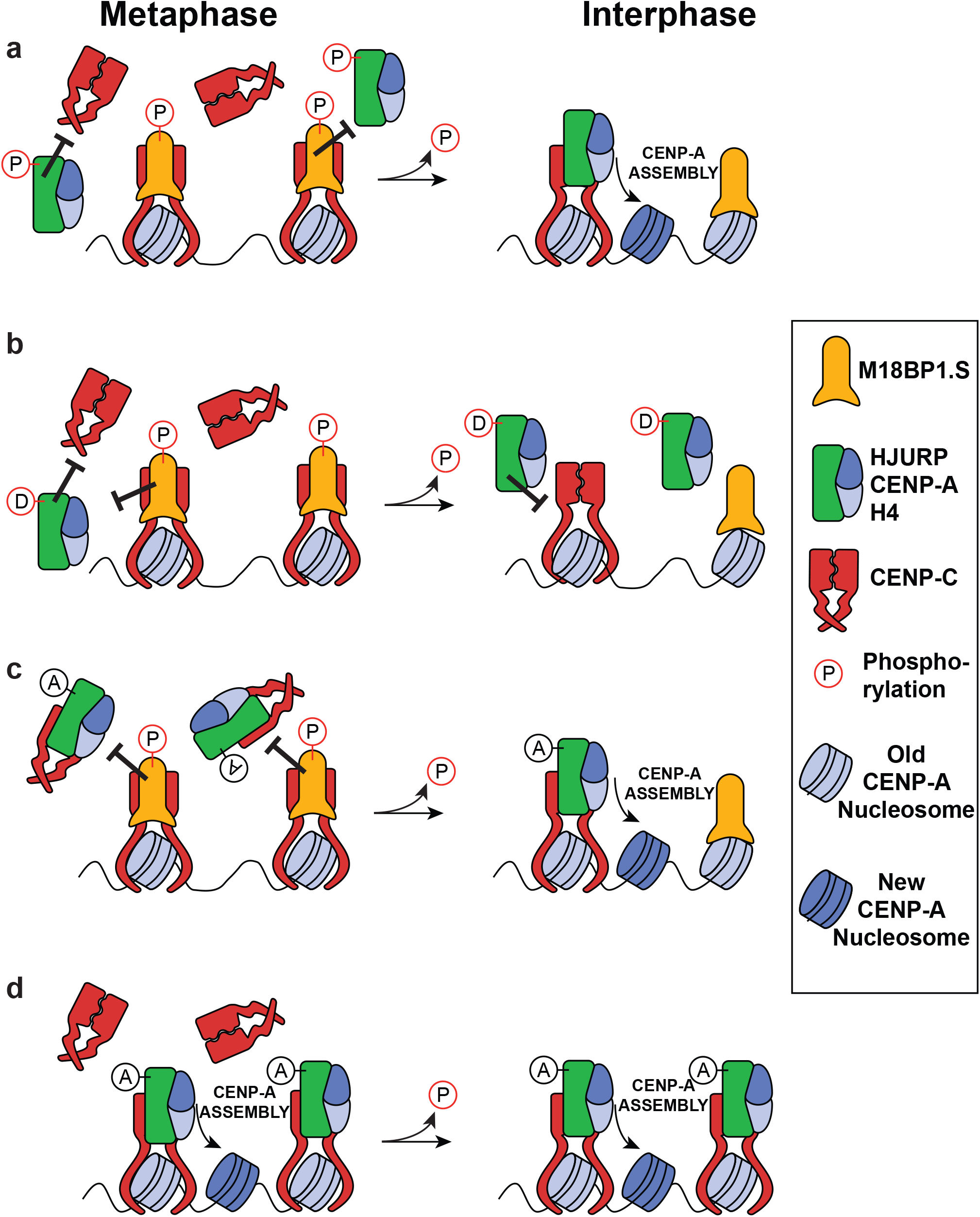
Model for dual regulation of CENP-A assembly through HJURP and M18BP1.S interaction with the CENP-C cupin domain. a) In metaphase, the targeting of the HJURP/CENP-A/H4 complex to the centromere is inhibited by phosphorylation of HJURP^S220^ preventing interaction with CENP-C. Phosphorylated M18BP1.S binds to the cupin domain of CENP-C providing a second inhibitory mechanism that prevents HJURP interaction in metaphase. Upon cycling into interphase, the inhibitory phosphorylation of HJURP^S220^ is removed and M18BP1.S dissociates from the CENP-C cupin domain allowing HJURP targeting to centromeres and new CENP-A assembly. b) The HJURP^S220D^ mutation mimics mitotic phosphorylation and blocks HJURP binding to CENP-C. This phosphomimetic HJURP is unable to dock onto the CENP-C cupin domain and thus cannot assemble new CENP-A in interphase c) The HJURP^S220A^ mutation that prevents mitotic phosphorylation of HJURP is able to bind to CENP-C in metaphase. M18BP1.S binding to CENP-C on chromatin prevents docking of the HJURP/CENP-A/H4 complex at centromeres. In interphase, M18BP1.S dissociates from CENP-C and the HJURP^S220A^/CENP-A/H4 complex binds to CENP-C and supports new CENP-A assembly. d) Dual perturbation by HJURP^S220A^ mutation and M18BP1.S depletion allows HJURP^S220A^/CENP-A/H4 to bind CENP-C and assemble new CENP-A in metaphase or interphase.

## DISCUSSION

In vertebrate cells, the assembly of new CENP-A nucleosomes is uncoupled from DNA replication and occurs when cells exit metaphase and transit into G1/interphase. We have identified two new regulatory mechanisms that control the timing of new CENP-A assembly during the cell cycle in *Xenopus laevis*. We find that CENP-A assembly is inhibited in metaphase by phosphorylation of the conserved serine 220 of the HJURP chaperone. Phosphorylation of HJURP^S220^ prevents HJURP interaction with CENP-C thereby inhibiting the delivery of soluble CENP-A to the centromere (Fig. 6a). The exit from metaphase and dephosphorylation of HJURP^S220^ reverses this inhibition. The phosphomimetic mutant HJURP^S220D^ blocks HJURP binding to CENP-C in interphase and prevents CENP-A assembly (Fig. 6b). Mutation of HJURP^S220^ to alanine allows HJURP to bind to CENP-C in solution but this interaction is not sufficient to localize HJURP to chromatin or cause new CENP-A assembly (Fig. 6c). We find that the M18BP1 isoform M18BP1.S provides a second inhibitory mechanism by binding directly to CENP-C and competing for HJURP binding in metaphase. Upon release from metaphase, M18BP1.S dissociates from CENP-C allowing HJURP to bind CENP-C and deliver CENP-A/H4 to centromeres (Fig. 6a). Breaking these two inhibitory mechanisms through mutation of HJURP^S220^ and removal of M18BP1.S causes premature CENP-A assembly in metaphase (Fig. 6d).

The CENP-C cupin domain provides an evolutionarily conserved protein binding interface for the regulation of CENP-A nucleosome assembly. The C-terminal cupin domain of CENP-C homodimerizes (Chik et al., 2019; Medina-Pritchard et al., 2020; Cohen et al., 2008; Sugimoto et al., 1997) and binds to the HJURP chaperone in vertebrates (French et al., 2017; Tachiwana et al., 2015) and to the CENP-A chaperone Cal1 in *Drosophila melanogaster* (Medina-Pritchard *et al*., *2020; Schittenhelm et al*., *2010). Dimerization of the cupin domain is required for HJURP interaction with CENP-C in Xenopus laevis* (Fig. 1c) and for Cal1 interaction with *D. melanogaster* CENP-C (Medina-Pritchard et al., 2020). We have previously shown that M18BP1.S and CENP-C interact through the C-terminus of CENP-C and this interaction occurs in metaphase (Moree et al., 2011; French and Straight, 2019). The Mis18 complex is known to play a stimulatory role for new CENP-A assembly in interphase. Here we find that the metaphase binding of M18BP1.S to the CENP-C cupin domain uncovers an additional role for M18BP1.S as a competitive inhibitor of HJURP binding and a negative regulator of new CENP-A assembly. *Xenopus laevis* is allotetraploid and expresses two homologues of M18BP1 from the short and long chromosomes (Session et al., 2016). The M18BP1.S and M18BP1.L proteins differ in their functions and timing of localization to centromeres such that only the M18BP1.S homeologue binds to centromeres in metaphase, depends upon CENP-C for the interaction (Moree et al., 2011), and acts as an inhibitor of HJURP binding (Fig. 4 and Extended Data Fig. 3). It will be interesting to determine whether organisms that express a single M18BP1 homolog use M18BP1 as both an inhibitor and activator of CENP-A assembly.

Mitotic kinases play a central role in controlling the timing of new CENP-A assembly (Srivastava et al., 2018). In particular, CDK phosphorylation of HJURP controls its localization to centromeres and its association with the Mis18 complex (Müller et al., 2014; Wang et al., 2014; Stankovic et al., 2017). Here we identify a new mitotic phosphorylation dependent regulatory mechanism that governs the activation of new CENP-A assembly as cells exit metaphase. HJURP Serine 220 phosphorylation blocks the interaction of HJURP with CENP-C so that soluble CENP-A cannot be delivered to the centromere for new CENP-A assembly. Phosphorylation profiling of human HJURP has shown that the conserved S185 residue is phosphorylated in M-phase and quickly dephosphorylated in early G1 (Stankovic et al., 2017), suggesting that *Xenopus* and humans may share this regulatory mechanism. In parallel, CDK activity dissociates the M18BP1 protein from the Mis18α/Misβ subunits of the complex (McKinley and Cheeseman, 2014; Pan et al., 2017; Spiller et al., 2017; French and Straight, 2019) while promoting the M18BP1.S interaction with CENP-C (French and Straight, 2019), thereby preventing HJURP access to the centromere. Artificial tethering of HJURP to chromatin has been shown to be sufficient for causing CENP-A deposition into chromatin (Hori et al., 2013; Barnhart et al., 2011; Shono et al., 2015; French et al., 2017) thus this dual inhibitory mechanism may prevent spurious CENP-A assembly if one pathway is compromised.

One interesting observation from this work is that despite the HJURP^S220A^ mutant’s ability to bind to soluble CENP-C in metaphase it is unable to displace M18BP1.S from CENP-C bound to the centromere. We have previously shown that M18BP1.S can bind both soluble and chromatin associated CENP-C in metaphase and depends on phosphorylation of M18BP1.S^T166^ and the conserved SANTA domain (Moree et al., 2011; French and Straight, 2019). Thus one speculative model is that CENP-C bound to CENP-A nucleosomes provides a preferred or higher affinity binding site for M18BP1.S than does soluble CENP-C. In humans, in addition to M18BP1 interaction with CENP-C, the Mis18β subunit of the Mis18 complex also interacts with CENP-C through the C-terminal region of the CENP-C protein (Stellfox et al., 2016). It will be interesting to explore whether M18BP1.S uses additional chromatin dependent interactions with metaphase centromeres to increase its local affinity and prevent HJURP binding to centromeres until exit into interphase.

## MATERIALS AND METHODS

### Experimental Model and Subject Details

Mature *Xenopus laevis* females (Nasco, LM00535MX) were housed and maintained in the Stanford Aquatic Facility staffed by the Veterinary Service Center. For ovulation, frogs were primed 2-14 days before ovulation by subcutaneous injection of 50 U pregnant mare serum gonadotropin (PMSG; Sigma) at the dorsal lymph sac, then induced 18 hours before ovulation with 500 U human chorionic gonadotropin (hCG; Chorulon). During ovulation, frogs were housed individually in 2 L MMR buffer (6 mM Na-HEPES, pH 7.8, 0.1 mM EDTA, 100 mM NaCl, 2 mM KCl, 1 mM MgCl2, and 2 mM CaCl2) at 17°C. Animal work was carried out in accordance with the guidelines of the Stanford University Administrative Panel on Laboratory Animal Care (APLAC).

### Protein Purification

*X. laevis* CENP-A-H4-Myc tetramer were expressed as soluble tetramers and purified as an adaptation of Guse et al., 2012. Briefly, CENP-A/H4 was expressed from a pST39 vector in BL21 (DE3) Codon Plus RIPL Escherichia coli (Agilent Technologies, 230280), grown in 2xYT medium (20 g/liter tryptone, 10 g/liter yeast extract, and 5 g/liter NaCl), and induced for 4-5 hours with 0.25 mM IPTG at an OD600 of 0.5 – 0.6 at room temperature. Bacteria were harvested and resuspended in lysis buffer (20 mM potassium phosphate, pH 6.8, 1M NaCl, 5 mM 2-mercaptoethanol, 1 mM PMSF, 1 mM benzamidine, 0.05% NP-40, and 0.2 mg/ml lysozyme), homogenized using an EmulsiFlex-C5 (Avestin, Inc.) followed by 1 × 30 second sonication. Lysate was run through a hydroxyapatite (HA) column (type II 20 mM HA; Bio-Rad Laboratories) pre-equilibrated with 20 mM potassium phosphate, pH 6.8. After washing with 6 column volumes (20 mM potassium phosphate, pH 6.8, 1 M NaCl, and 5 mM 2-mercaptoethanol) tetramer was eluted with HA buffer containing 3.5 M NaCl. The elution was dialyzed into 10 mM Tris-HCl, pH 7.4, 0.75 M NaCl, 10 mM 2-mercaptoethanol, and 0.5 mM EDTA. The dialyzed protein was passed through a 1 ml HiTrap SP FastFlow column, washed, and tetramer was eluted over a 20–column volume gradient into dialysis buffer containing 2 M NaCl. Fractions containing concentrated tetramer were pooled, aliquoted, frozen in liquid nitrogen, and stored at −80°C.

MBP-M18BP1.S-161-580 and MBP-M18BP1.S-161-580 SANTA were independently expressed from modified pMal-c2x plasmids (NEB) in BL21 (DE3) Codon Plus RIPL E. coli. For each purification, two 2L cultures were grown in 2xYT medium to OD600 of 0.6 at 37°C and then induced with 0.5 mM IPTG for 4 hours at 21°C. Bacteria were lysed in MBP lysis buffer (10 mM sodium phosphate pH 7.4, 500 mM NaCl, 2.7 mM KCl, 1 mM EDTA, 1 mM dithiothreitol (DTT), 1 mM phenylmethylsulfonyl fluoride (PMSF), 1 mM benzamidine hydrochloride, 10 μg/mL LPC [leupeptin, pepstatin A, chymostatin], 0.2 mg/mL lysozyme) by three passes through an EmulsiFlex-C5 (Avestin, Inc.) and then clarified by centrifugation at 40,000 rpm in a Type 45ti rotor (Beckman) for 1.5 hours at 4°C. Clarified lysates were flowed over 2 mL amylose resin (NEB) equilibrated with MBP lysis buffer to capture MBP-tagged protein. Resin was subsequently washed with 30 column volumes lysis buffer. Bound protein was then washed with 10 column volumes MBP wash buffer (20 mM Na-HEPES pH 7.7, 150 mM NaCl, 5 mM 2-mercaptoethanol) and eluted with MBP wash buffer containing 10 mM maltose. Protein was dialyzed against protein storage buffer (20 mM Na-HEPES, pH 7.7, 150 mM NaCl, 5 mM 2-mercaptoethanol, 10% glycerol), concentrated to 1 mL by ultracentrifugation (Amicon Ultra 15 MWCO 30k; Millipore), and stored in 50 μL aliquots at −80°C.

### *In Vitro* Translation

*In vitro* translated (IVT) proteins were generated in rabbit reticulocyte lysate using the SP6 TNT Quick-Coupled Transcription/Translation kit (Promega, L2080) according to the manufacturer’s instructions.

### *X. laevis* Egg Extracts

CSF-arrested (metaphase) *Xenopus* egg extracts were prepared as described previously (Desai et al., 1999; Guse et al., 2012). In brief, eggs were washed with MMR buffer (6 mM Na-HEPES, pH 7.8, 0.1 mM EDTA, 100 mM NaCl, 2 mM KCl, 1 mM MgCl2, and 2 mM CaCl2) and then de-jellied in MMR + 2% L-cysteine. De-jellied eggs were washed in CSF-XB buffer (10 mM K-HEPES, pH 7.7, 100 mM KCl, 50 mM sucrose, 2 mM MgCl2, 0.1 mM CaCl2, 5 mM EGTA), followed by CSF-XB containing 10 mg/ml LPC [leupeptin/pepstatin A/chymostatin]. Eggs were packed in a 13×51-mm polyallomer tube (Beckman Coulter, 326819) by a low-speed spin in a table top clinical centrifuge for 45 s. After removal of buffer, packed eggs were centrifuged in a SW55Ti rotor (Beckman Coulter) for 15 min at 10,000 rpm. The soluble cytoplasm was removed by side puncture with a 18G needle and supplemented with 50 mM sucrose, 10 mg/ml LPC, 10 mg/ml cytochalasin D, and energy mix (7.5 mM creatine phosphate, 1 mM ATP, and 1 mM MgCl2).

### Immunodepletions

Immunodepletion of CENP-C and M18BP1 from Xenopus extract was performed essentially as described ((Moree et al., 2011)). Protein A Dynabeads (Invitrogen, 10001D) were washed in TBSTx (50 mM Tris-HCl, pH 7.4, 150 mM NaCl, 0.1% Triton X-100) or CSF-XBT and then coupled to affinity purified antibody for 30-60 minutes at 4°C. For 100 μL extract, 5 μg anti-xCENP-C antibody (rabbit, raised and purified against xCENP-C^207–296^ (Milks et al., 2009)), or 10 μg anti-xM18BP1 (rabbit, raised against GSTxM18BP1.L amino acids 161–415 and purified against xM18BP1.S 161–375 (Moree et al., 2011) were used. An equivalent amount of whole rabbit IgG (Jackson ImmunoResearch Laboratories, Inc.) was used for mock depletions. Antibody-coupled beads were washed three times in TBSTx or CSF-XBT and then re-suspended in extract. Depletions were rotated at 4°C for 1 hour. Beads were removed from extract by three rounds of 5-minute magnetizations.

### CENP-A Assembly Assays

CENP-A assembly assays on sperm chromatin or reconstituted chromatin were performed as described previously (Moree et al., 2011; Westhorpe et al., 2015). RNAs used to express protein in egg extract were prepared with the SP6 mMessage mMachine kit (Life Technologies, AM1340). NotI linearized pCS2+ plasmids were used in the transcription reaction as per manufacturer’s instructions, using double the amount of DNA. RNA was purified using RNeasy mini columns (QIAGEN). RNA was typically concentrated to at least 1 μg/uL using a SpeedVac (Savant) to avoid excessive dilution of the extract. For sperm assembly assays, 50 μL assembly reactions contained 1.5 μL xFLAG-HJURP IVT protein and/or 1.5 μL xFLAG-CENP-C IVT protein, and 25 ng/uL V5-xCENP-A RNA. CSF-arrested, CENP-C-depleted extract (prepared as above) was supplemented with IVT protein and RNA and incubated at 16-18°C for 30 minutes to permit RNA translation. Cycloheximide was added to 0.1 mg/mL to terminate translation. Reactions were then released to interphase by addition of 750 uM CaCl2 and supplemented with demembranated sperm chromatin to 3,000/uL. After incubation for 75 minutes at 16-18°C, assembly reactions were diluted in 1 mL dilution buffer (BRB-80 [80 mM K-PIPES, pH 6.8, 1 Mm MgCl2, 1 mM EGTA], 0.5% Triton X-100, 30% glycerol) containing 150 mM KCl and incubated on ice for 5 minutes. These samples were then fixed by addition of 1 mL dilution buffer containing 4% formaldehyde and spun through cushions of 40% glycerol in BRB-80 onto poly-L-lysine-coated coverslips to be processed for immunofluorescence. Quantification of CENP-A loading (centromeric V5-CENP-A signal) was performed as described below.

### Immunofluorescence Preparation

Coverslips were first blocked in antibody dilution buffer (AbDil; 20 mM Tris-HCl, pH 7.4, 150 mM NaCl with 0.1% Triton X-100, and 2% bovine serum albumin) for 30 minutes. Coverslips were exposed to primary antibody diluted in AbDil for 15-30 minutes, washed three times with AbDil, and then exposed to AlexaFluor-conjugated secondary antibodies (Life Technologies; Jackson ImmunoResearch Laboratories, Inc.). When required, coverslips were blocked with 1 mg/mL whole rabbit or whole mouse IgG in AbDil followed by staining with directly-conjugated primary antibodies. Sperm nuclei were stained with 10 μg/mL Hoechst 33258 in AbDil for DNA. Coverslips were mounted in 90% glycerol, 10 mM Tris-HCl, pH 8.8, 0.5% p-phenylenediamine, sealed to a slide with clear nail polish and stored at −20°C. Primary antibodies used were: 1 mg/mL anti-FLAG (Sigma, F7425 [rabbit] and F1804 [mouse]), 0.5 mg/mL mouse anti-Myc (EMD Millipore, 4A6 [mouse]), 1 mg/mL rabbit anti-Myc (EMD Millipore, 06-549 [rabbit]), 1:500 mouse anti-V5 (Thermo Fisher, R960-25), 1 mg/mL rabbit-anti-xCENP-A. Secondary antibodies used were AlexaFluor 488-, 568-, and 647-conjugated goat anti-mouse or anti-rabbit (Life Technologies), and AlexaFluor 647-conjugated donkey anti-rabbit (Jackson ImmunoResearch Laboratories, Inc.). All secondaries were used at 1 mg/mL.

### Image Acquisition and Processing

Imaging was performed on an IX70 Olympus microscope with a DeltaVision system (Applied Precision), a Sedat quad-pass filter set (Semrock #DA/FI/TR/Cy5-4×4M-C-000), monochromatic solid-state illuminators (Lumencor). Device control utilized softWoRx 4.1.0 software (Applied Precision). Images were acquired using a 60x 1.4 NA Plan Apochromat oil immersion lens (Olympus) with a charge-coupled device camera (CoolSNAP HQ2; Photometrics) and digitized to 12 bits. Z-sections were taken at 0.2-μm intervals. Displayed images of sperm nuclei are maximum intensity projections of deconvolved z-stacks. Image analysis of sperm chromatin (Moree et al., 2011) was performed using custom software as previously described. For sperm, at least 250 centromeres among at least 16 nuclei were counted in each condition in each experiment.

### Immunoblotting

Samples were resolved by SDS-PAGE and transferred onto polyvinylidene fluoride membrane (Bio-Rad Laboratories). For immunoblotting CENP-C, samples were transferred in CAPS transfer buffer (10 mM 3- (cyclohexylamino)-1-propanesulfonic acid, pH 11.3, 0.1% SDS, 20% methanol). All other samples were transferred in Tris-glycine transfer buffer (20 mM Tris-HCl, 200 mM glycine, 20% methanol). Samples containing 1.5 μL Xenopus egg extract or 1 – 1.5 μL IVT protein were loaded per lane. Membranes were blocked in 3% milk in 1X TBST (20 mM Tris base pH 7.4, 150 mM NaCl, 2.7 mM KCl, 0.2% Tween 20), probed with primary antibody, and detected using horseradish peroxidase-conjugated goat-a-mouse or goat-a-rabbit secondary antibodies followed by chemiluminescence (Thermo Scientific). Chemiluminescent membranes were exposed to X-ray film for 5–180 s and developed with a Konica SRX-101A film processor. Primary antibodies used were: 1 μg/mL mouse-anti-FLAG, 1 μg/mL mouse-anti-Myc, 1:2500 mouse anti-V5, 10 μg/mL rabbit-antixM18BP1, 2 μg/mL rabbit-anti-xHJURP (raised against GST-xHJURP^42-194^ and purified against His-xHJURP^42-194^), and 5 μg/mL rabbit-anti-xCENP-C. All secondary antibodies were used at 0.5 μg/mL.

### Immunoprecipitation

To assay association of endogenous HJURP with CENP-C, 100 μL CSF-arrested or interphase extract reactions were added to 10 μL Protein A Dynabeads (Invitrogen) coupled to 5 μg rabbit anti-xCENP-C antibody. To assay association of endogenous HJURP with Myc-CENP-C truncations, 100 μL reactions containing 10 μL Myc-xCENP-C truncations in interphase CENP-C-depleted extract were added to 10 μL Protein A Dynabeads (Invitrogen) coupled to 2.5 μg mouse anti-Myc antibody. To assay association of endogenous HJURP with FLAG-CENP-C mutants, 100 μL reactions containing 10 μL FLAG-xCENP-C mutants in interphase CENP-C-depleted extract were added to 10 μL Protein A Dynabeads (Invitrogen) coupled to 2.5 μg mouse anti-FLAG antibody. To assay association of endogenous CENP-C with FLAG-HJURP truncations or mutants, 100 μL reactions containing 10 μL FLAG-xHJURP truncations or mutants in CSF-arrested or interphase extract were added to 10 μL Protein A Dynabeads (Invitrogen) coupled to 2.5 μg mouse anti-FLAG antibody. Following rotation for 1 hour at 4°C, beads were collected by magnetization and washed four times in TBSTx. Immunoprecipitates were eluted in sample buffer (50 mM Tris, pH 6.8, 15 mM EDTA, 1 M b-mercaptoethanol, 3.3% SDS, 10% glycerol, and 1 mg/ml Bromophenol blue), resolved by SDS-PAGE, and immunoblotted as described above.

To assay direct binding of HJURP and CENP-C, FLAG-HJURP or FLAG-HURP mutants and Myc-CENP-C were translated *in vitro* as described above. 50 μL binding reactions containing 10 μL FLAG-HJURP and 10 μL Myc-CENP-C in IVT binding buffer (50 mM Tris-HCl, pH 8.0, 50 mM NaCl, 0.25 mM EDTA, 0.05% Triton X-100, and 2% BSA) were incubated at 4°C for 1 h. Bound material was recovered by subsequent incubation of binding reactions with 10 mL Protein A Dynabeads (Invitrogen) coupled to 2.5 mg mouse anti-FLAG at 4°C for 1 h. Beads were then washed three times with TBSTx and bound material eluted with sample buffer. Samples were resolved by SDS-PAGE and bound material visualized by chemiluminescence.

### Protein Alignment

Protein sequences homologous to *X. laevis* HJURP were identified using NCBI PSI-BLAST with an input query of amino acids 205 - 225. Multiple sequence alignments were performed with MAFFT (version 7.305b) (Katoh et al., 2002) using default parameters.

### Quantification and Statistical Analysis

Image analysis of *X. laevis* sperm centromeres was performed using custom software as previously described ((Moree et al., 2011)). To identify centromere regions, images were normalized by median filtering the image and then dividing the original image by the filtered intensity value. Centromere masks were then generated by thresholding the channel with the indicated centromere marker (endogenous CENP-A) followed by size filtering to remove regions larger or smaller than centromeres. Masks were then manually inspected to remove any remaining errant regions. Mean pixel intensity per centromere region was then measured for the channels of interest from maximum intensity projections. Reported values represent the average of this quantity across all centromere regions per condition, normalized to the indicated positive control. For sperm, at least 250 centromeres among at least 16 nuclei were counted in each condition in each experiment.

### Reagents Used in This Study

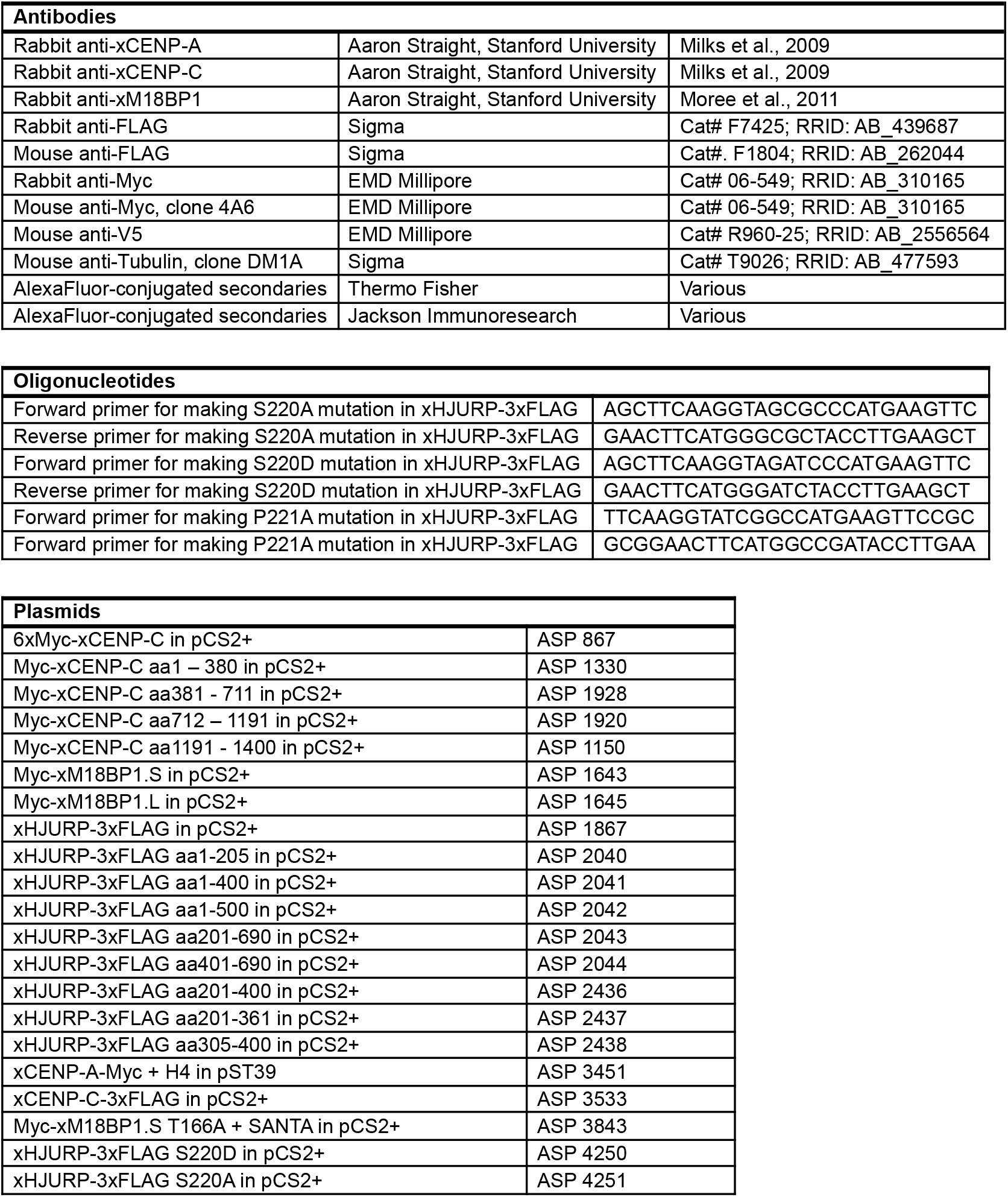
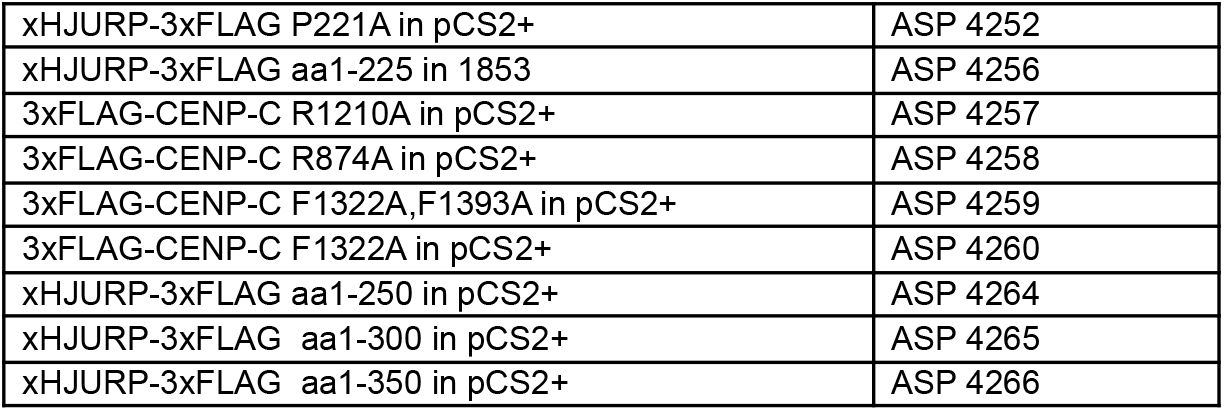

## ACKNOWLEDGEMENTS

We are grateful to Rachel Brown, Bradley French and members of the Straight Laboratory for feedback on the manuscript. We thank Bradley French for providing MBP-M18BP1.S-161-580 protein. This work was supported by NIH R01 GM074728 to AFS and the Charles Yanofsky Fellowship to JFS.

## AUTHOR CONTRIBUTIONS

Julio Flores Servin: Conceptualization, Methodology, Validation, Investigation, Resources, Data Curation, Writing - Original Draft, Writing - Review and Editing, Visualization

Aaron F. Straight: Conceptualization, Methodology, Validation, Resources, Writing - Original Draft, Writing - Review and Editing, Visualization, Supervision, Project Administration, Funding Acquisition

## ABBREVIATIONS

CENP: centromere protein
HJURP: Holliday junction recognition protein
M18BP1: Mis18 binding protein 1
CDK1: cyclin dependent kinase 1

---

**Supplementary Figure 1.**
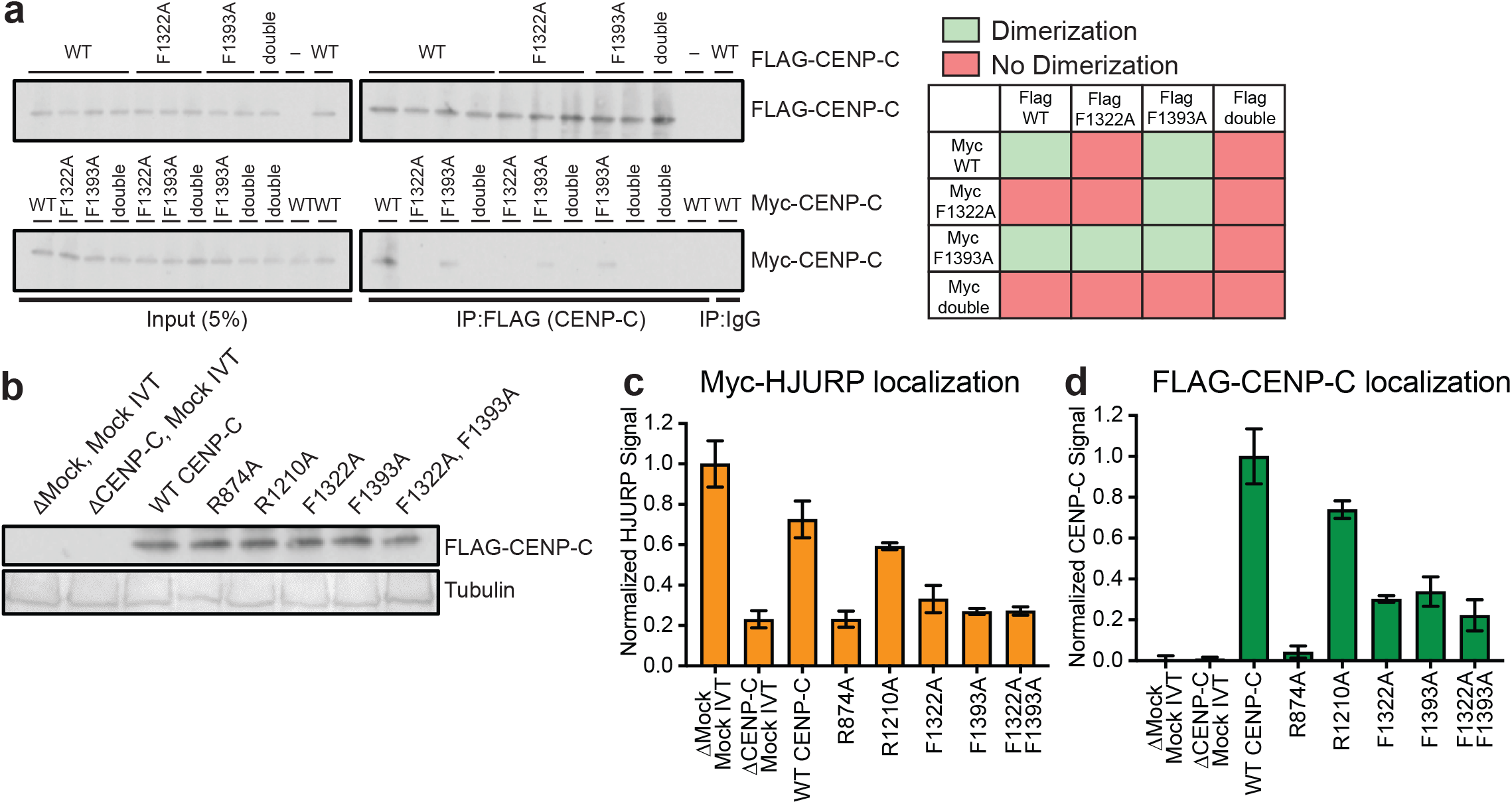
Analysis of effect of CENP-C mutations on HJURP and CENP-C activity. a) Mutations in the CENP-C cupin domain inhibit CENP-C dimerization in *Xenopus* egg extract. Interphase extract was supplemented with the specified FLAG-CENP-C and Myc-CENP-C mutants. The F1322A/F1393A mutant is labeled double. Co-immunoprecipitation of Myc-CENP-C (bottom panels) was assayed by anti-Myc immunoblotting following FLAG-CENP-C precipitation (top panels). Mock precipitations using scrambled DNA translated *in-vitro* (-IVT) or whole mouse IgG are indicated. The left panels contain 5% of the input material and the right panel contain the immunoprecipitations. A color coded matrix showing combinations of mutants that block (red) or allow (green) dimerization is displayed on the right. b) Levels of *in vitro* translated mutant CENP-C proteins after addition to *X. laevis* egg extracts used in **Fig. 1e,f**. Tubulin is shown as a loading control, the mutant or wild type protein used is indicated above each lane. c) HJURP centromere localization is affected when CENP-C depleted interphase extracts are rescued with FLAG-CENP-C mutants. Quantification of Myc-HJURP centromere localization on sperm chromatin in CENP-C depleted interphase extracts rescued with specified CENP-C mutant. Values are normalized to mock depletion and mock IVT centromere signal in interphase extract. Plot shows mean Myc-HJURP signal on sperm chromatin ± SEM of at least three experiments. d) Mutations to the CENP-A binding region and the cupin domain of CENP-C negatively affect centromere localization. Quantification of FLAG-CENP-C centromere localization on sperm chromatin in CENP-C depleted interphase extracts rescued with specified CENP-C mutant. Values are normalized to CENP-C depletion and wild type CENP-C IVT add-back signal in interphase extract. Plot shows mean FLAG-CENP-C signal on sperm chromatin ± SEM of at least three experiments.

**Supplementary Figure 2.**
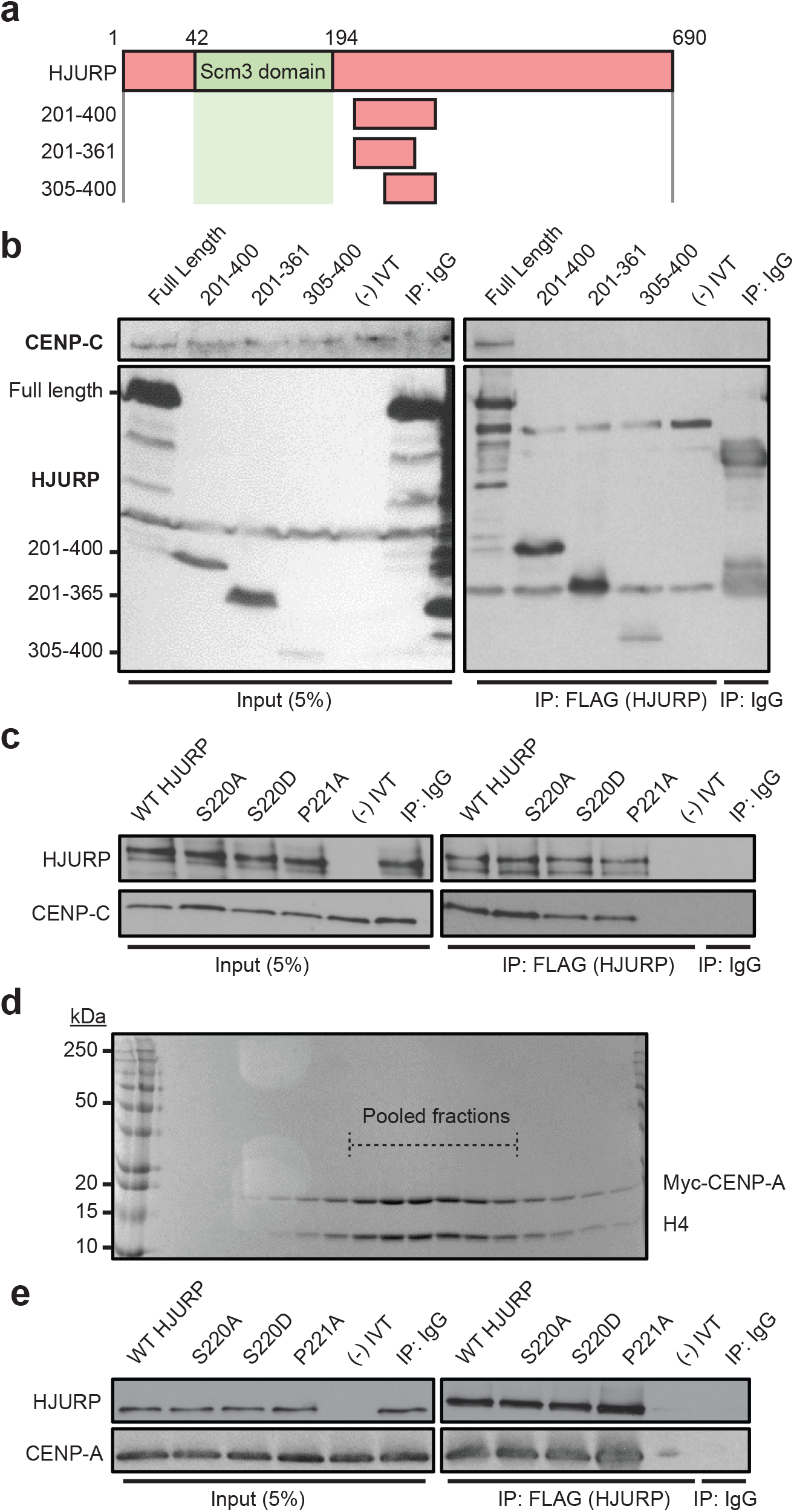
Mutations on the conserved N-terminal SP residues of HJURP do not affect CENP-C and CENP-A binding *in vitro*. a) Schematic representation of HJURP truncations used to identify the CENP-C interacting region in **(b)**. b) Fragments of HJURP C-terminal to amino acid residue 201 do not bind CENP-C in interphase extracts. Interphase extract was supplemented with specified FLAG-HJURP truncations. Co-immunoprecipitation of endogenous CENP-C was assayed by anti-CENP-C immunoblotting following immunoprecipitation of FLAG-HJURP. Mock precipitations using scrambled DNA translated *in-vitro* (-IVT) or whole mouse IgG served as negative controls. c) S220 and P221 mutations on HJURP do not affect CENP-C binding *in vitro. In vitro* interaction assay was performed using *in vitro* translated HJURP mutant and CENP-C. Co-immunoprecipitation of CENP-C was assayed by anti-CENP-C immunoblotting following FLAG-HJURP mutant precipitation. Mock precipitations using scrambled DNA translated *in-vitro* (-IVT) or whole mouse IgG served as negative controls. d) Purification of recombinant Myc-CENP-A/H4 heterodimer. Coomasie stained SDS-PAGE gel of S-column fractions of purified Myc-CENP-A/H4 heterodimer. Dotted lines highlight pooled fractions. e) Binding of CENP-A/H4 to HJURP is not affected by S220 or P221 mutations. *In vitro* interaction assay was performed using the indicated *in vitro* translated HJURP mutant and recombinant purified *Xenopus* CENP-A/H4. Co-immunoprecipitation of endogenous CENP-A was assayed by anti-CENP-A immunoblotting following FLAG-HJURP precipitation. Mock precipitations using scrambled DNA translated *in-vitro* (-IVT) or whole mouse IgG served as negative controls.

**Supplementary Figure 3.**
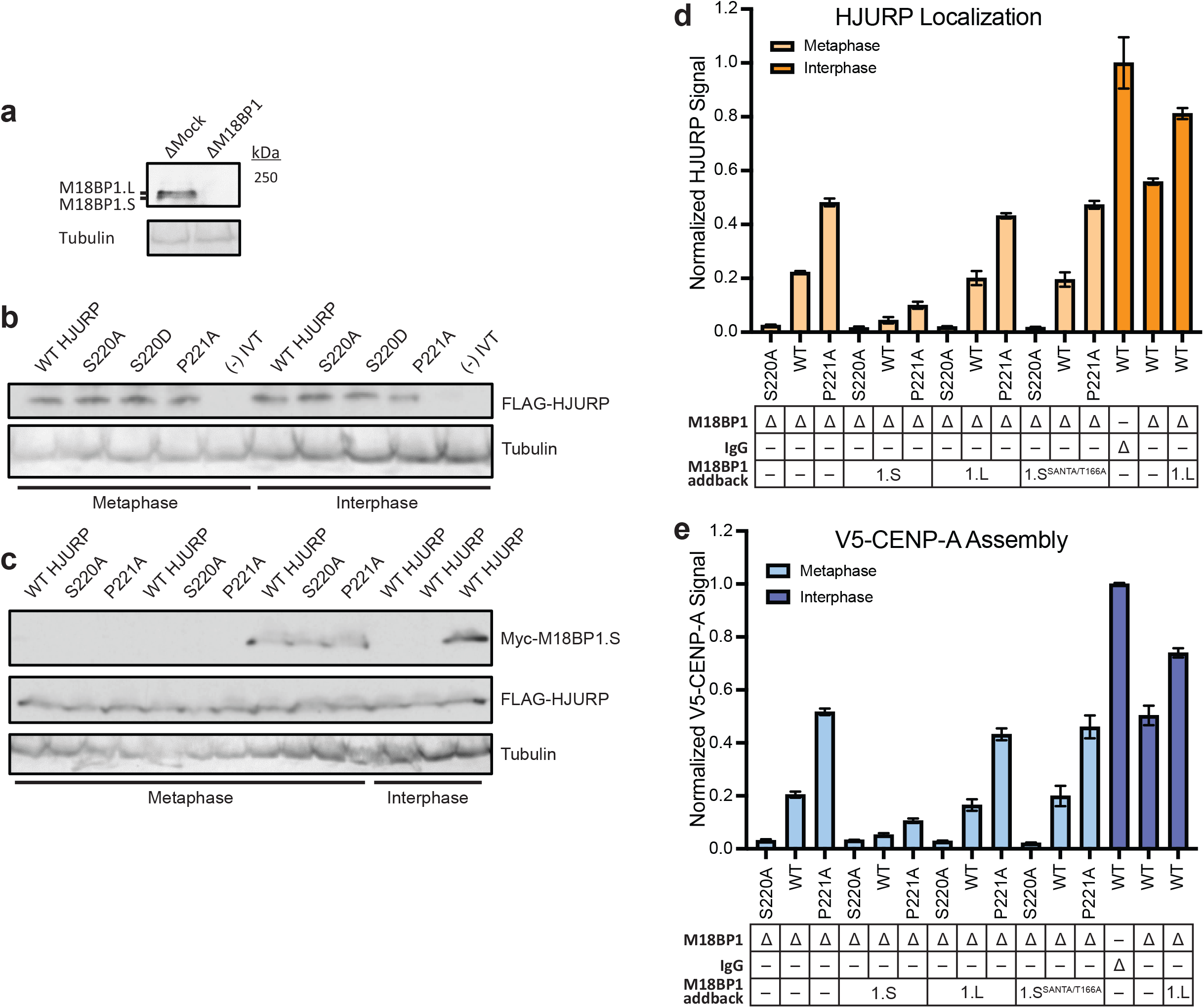
M18BP1.S mutants that cannot localize in metaphase and M18BP1.L do not prevent premature HJURP localization and CENP-A assembly. a) Representative western blot of M18BP1 depleted extract samples used in **Fig. 4a,b,c**. Depletion (Δ) is indicated above on each column. A tubulin western blot is included as a loading control. b) *In vitro* translated mutant HJURP proteins added into extracts for experiments in **Fig. 3a,b,c**. Samples were taken from egg extracts and western blotted for FLAG(HJURP) and Tubulin as a loading control. c) *In vitro* translated mutant HJURP and M18BP1.S proteins added into extracts for experiments in **Fig. 4a,b,c**. Samples were taken from egg extracts and western blotted for FLAG(HJURP), Myc(M18BP1) and Tubulin as a loading control. d) The M18BP1.L protein that does not bind CENP-C in metaphase or an M18BP1.S mutant (M18BP1.S^SANTA/T166A^) that cannot bind CENP-C fail to compete for HJURP localization at centromeres in metaphase extracts. Quantification of FLAG-HJURP centromere localization on sperm chromatin in M18BP1 depleted metaphase and interphase extracts. Values are normalized to wild type FLAG-HJURP centromere signal in mock depleted interphase extract. Bottom rows indicate depletion (Δ) status (M18BP1 or IgG antibody) and M18BP1 add-back (1.S, 1.L, or 1.S^SANTA/T166A^) for each condition. Plot shows mean FLAG-HJURP signal on sperm chromatin ± SEM of at least three experiments. e) Metaphase CENP-A assembly is not affected by addition of M18BP1.L and M18BP1.S^SANTA/T166A^. Quantification of V5-CENP-A assembly on sperm chromatin in M18BP1 depleted metaphase and interphase extracts. Values are normalized to V5-CENP-A assembly signal in wild type HJURP condition on mock depleted interphase extracts. Bottom rows indicate depletion (Δ) status (M18BP1 or IgG antibody) and M18BP1 add-back (1.S, 1.L, or 1.S^SANTA/T166A^) for each condition. Plot shows mean V5-CENP-A assembly signal on sperm chromatin ± SEM of at least three experiments.

**Supplementary Figure 4.**
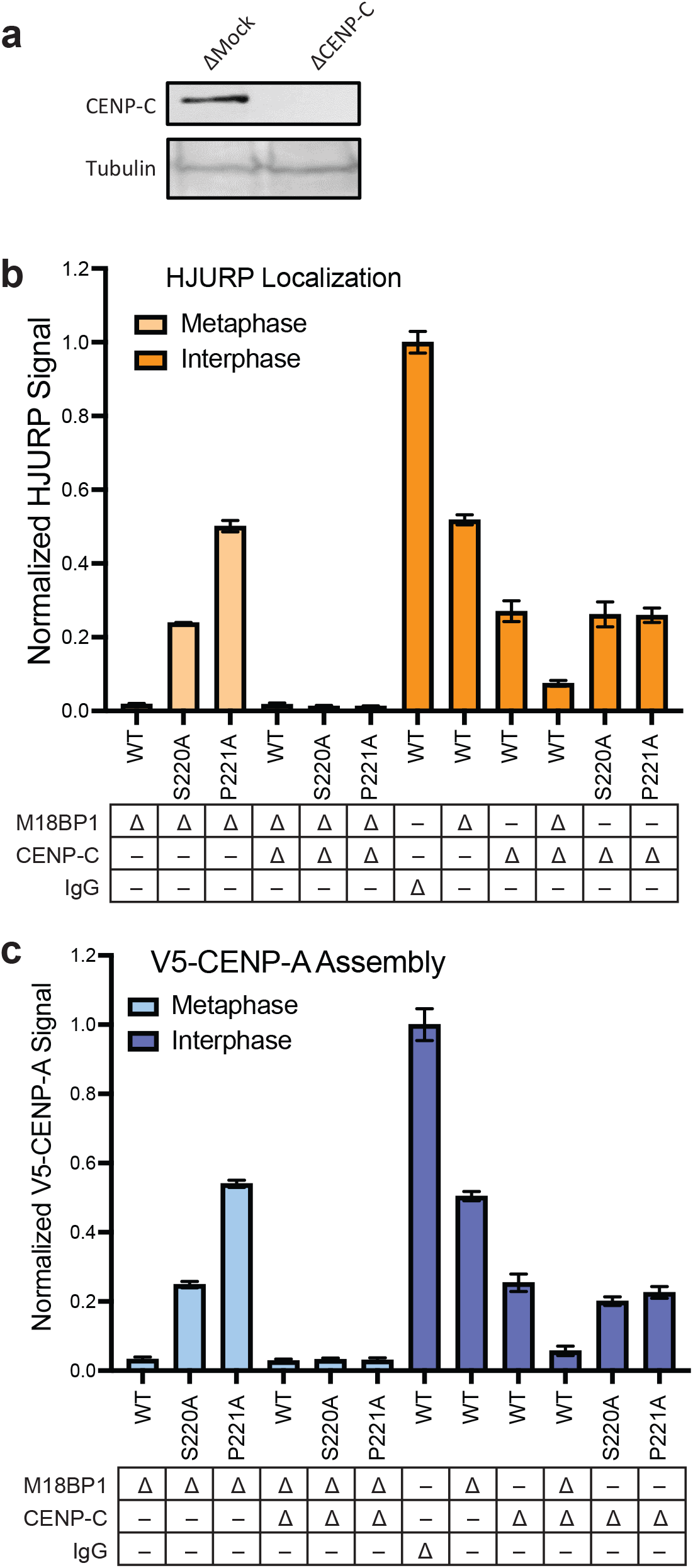
Metaphase HJURP centromere localization and CENP-A assembly require CENP-C. a) Representative western blot of CENP-C depleted extract samples used in **(b)** and **(c)**. Depletion (Δ) is indicated above each column. A tubulin western blot is included as a loading control. b) Dual depletion of CENP-C and M18BP1 in metaphase prevents localization of HJURP mutants. Quantification of FLAG-HJURP centromere localization on sperm chromatin in M18BP1 and/or CENP-C depleted metaphase and interphase extracts. Values are normalized to wild type FLAG-HJURP centromere signal in mock depleted interphase extract. Bottom rows indicate depletion (Δ) status (M18BP1, CENP-C, or IgG antibody) for each condition. Plot shows mean FLAG-HJURP signal on sperm chromatin ± SEM of at least three experiments. c) CENP-C is required for premature CENP-A assembly in metaphase driven by HJURP^S220A or P221A^ and M18BP1 depletion. Quantification of V5-CENP-A assembly on sperm chromatin in single depletion (M18BP1) interphase and metaphase extracts, and dual depleted (CENP-C and M18BP1) extracts. Values are normalized to V5-CENP-A assembly signal in wild type HJURP condition on mock depleted interphase extracts. Bottom rows indicate depletion (Δ) status (M18BP1, CENP-C, or IgG antibody) for each condition. Plot shows mean V5-CENP-A assembly signal on sperm chromatin ± SEM of at least three experiments.

